# Aurora B switches relative strength between kinetochore–microtubule attachment modes to promote error correction

**DOI:** 10.1101/455873

**Authors:** Harinath Doodhi, Taciana Kasciukovic, Lesley Clayton, Tomoyuki U. Tanaka

**Affiliations:** Centre for Gene Regulation and Expression, School of Life Sciences, University of Dundee, Dow street, Dundee DD1 5EH, UK

## Abstract

For proper chromosome segregation, sister kinetochores must interact with microtubules from opposite spindle poles; this is called bi-orientation. To establish bi-orientation prior to chromosome segregation, any aberrant kinetochore–microtubule interaction must be resolved (error correction) by Aurora B kinase that phosphorylates outer kinetochore components. Aurora B differentially regulates kinetochore attachment to the microtubule plus end and its lateral side (end-on and lateral attachment, respectively). However, it is still not fully understood how kinetochore–microtubule interactions are exchanged during error correction. Here we reconstituted the kinetochore–microtubule interface of budding yeast *in vitro* by attaching the Ndc80 complexes (Ndc80C) to nanobeads. These Ndc80C–nanobeads recapitulated *in vitro* the lateral and end-on attachments of authentic kinetochores, on dynamic microtubules loaded with the Dam1 complex. This *in vitro* assay enabled the direct comparison of lateral and end-on attachment strength and showed that Dam1 phosphorylation by Aurora B makes the end-on attachment weaker than the lateral attachment. We suggest that the Dam1 phosphorylation weakens interaction with the Ndc80 complex, disrupts the end-on attachment and promotes the exchange to a new lateral attachment, leading to error correction. Our study reveals a fundamental mechanism of error correction for establishment of bi-orientation.

## Introduction

For accurate and successful chromosome segregation, kinetochores must interact properly with spindle microtubules (MTs) (Tanaka, 2010). The kinetochore initially interacts with the lateral side of a single MT (lateral attachment) and then becomes tethered at the MT plus end (end-on attachment) as the MT shrinks (Rieder and Alexander, 1990; Tanaka et al., 2007; Tanaka et al., 2005). Subsequently sister kinetochores form end-on attachments to MTs extending from opposite spindle poles, establishing chromosome bi-orientation. If an aberrant kinetochore–MT attachment is formed, it must be resolved (error correction) by Aurora B kinase (Ipl1 in budding yeast), which phosphorylates kinetochore components and disrupts the end-on attachment (Hauf et al., 2003; Lampson et al., 2004; Tanaka et al., 2002). In budding yeast, the Dam1 complex (Dam1C) is the most important Aurora B substrate for error correction (Cheeseman et al., 2002) and phosphorylation of Ndc80 N-terminus also contributes to this process (Akiyoshi et al., 2009).

We have shown that the end-on attachment is weakened by the action of Aurora B, but the lateral attachment is impervious to Aurora B regulation, i.e. the end-on and lateral attachments are differentially regulated (Kalantzaki et al., 2015). This led us to propose the model that, during error correction, an end-on attachment is disrupted by the action of Aurora B (Figure 1A, step 1, 2) and subsequently replaced by lateral attachment to a different MT (step 3, 4); the lateral attachment is then converted to end-on attachment and, if this forms aberrantly, it must be resolved again by Aurora B (step 1), but if bi-orientation is formed, tension across sister kinetochores stabilizes end-on attachment (step 5) (Kalantzaki et al., 2015). Thus, the model suggests that differential regulation of end-on and lateral attachments promotes the exchange of kinetochore–MT interactions for error correction.

**Figure 1.**
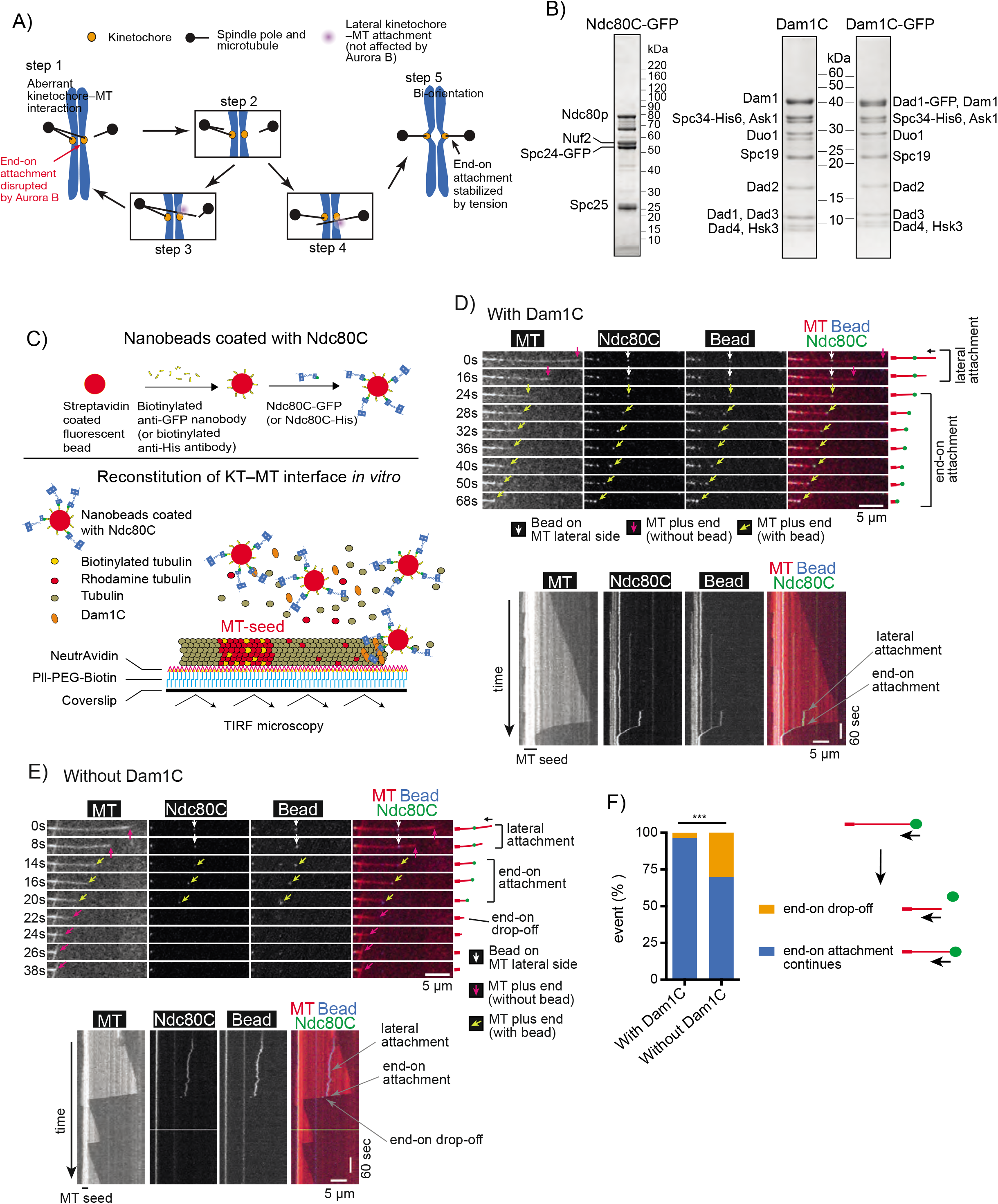
Behaviour of Ndc80C–nanobeads *in vitro* on dynamic MTs loaded with Dam1C. **A**) Diagram shows the model of an error correction process, proposed in (Kalantzaki et al., 2015). Each step is explained in the text. **B**) Coomassie Blue stained gels showing purified Ndc80C-GFP (with GFP at the C-terminus of Spc24), Dam1C and Dam1C-GFP (with GFP at the C-terminus of Dad1). **C**) Diagram shows that Ndc80C-GFP was attached to a streptavidin-coated nanobead through a biotinylated anti-GFP nanobody (top). Dynamic MTs were grown from stable MT seeds on coverslips in the presence of Dam1C and Ndc80C-GFP-coated nanobeads (Ndc80C–nanobeads) and observed by TIRF microscopy (bottom). **D**) Images in time sequence (top) show that an Ndc80C–nanobead formed MT lateral attachment (0–16 s time points) and end-on attachment (24–68 s) in the presence of Dam1C. Time 0 s was set arbitrarily. Scale bar, 5 μm. Also refer to diagrams (right). Kymograph was obtained for the same MT (bottom). Scale bars, 5 μm (horizontal) and 60 s (vertical). **E**) Images in time sequence (top) show that an Ndc80C–nanobead formed MT lateral attachment (0–8 s) and end-on attachment (14–20 s), and subsequently detached from the MT end (end-on drop-off; 22 s), in the absence of Dam1C. After the nanobead had detached from the MT end, it was not visible by TIRF microscopy because it was no longer close to the coverslip. Also refer to diagrams (right). Kymograph was obtained for the same MT (bottom). Scale bars are as in D). **F**) Graph shows the percentage of events; end-on drop-off and continuous end-on attachment, observed in the presence (n=82) and absence (n=30) of Dam1C. Difference between the two groups is significant (*p*=0.0003).

A number of crucial questions still remain regarding the exchange of kinetochore–MT interactions during error correction: For example, although studies in budding yeast cells suggested the differential regulation of end-on and lateral attachments by Aurora B (Kalantzaki et al., 2015), this has not yet been directly tested during the exchange of kinetochore–MT interactions as it is difficult to visualize in yeast cells. For exchange to occur efficiently, an end-on attachment to one MT must be disrupted and replaced by a lateral attachment to another MT. It is unknown how the relative strengths of lateral and end-on attachments are regulated by phosphorylation of kinetochore components by Aurora B kinase. Furthermore, although the Ndc80 complex (Ndc80C) and Dam1C are major outer kinetochore components comprising the kinetochore–MT interface (Jenni and Harrison, 2018; Kalantzaki et al., 2015; Lampert et al., 2010; Lampert et al., 2013; Sarangapani et al., 2013; Tien et al., 2010), it is unknown whether this interface, formed by the Ndc80C and Dam1C, sufficiently accounts for the differential regulation of lateral and end-on attachments by Aurora B kinase or whether any other factors are required for this regulation. To address these questions, we reconstituted the kinetochore–MT interface *in vitro* (i.e. in cell-free system) using recombinant Dam1C and Ndc80C and also with native kinetochore particles purified from budding yeast cells.

## Results

### Reconstitution of kinetochore–MT interface *in vitro* using defined outer kinetochore components

We aimed to reconstitute the kinetochore–MT interface *in vitro*, using nanobeads and recombinant Ndc80C and Dam1C. The diameter of the nanobead was about 100 nm, which is somewhat larger than the reported size of the inner kinetochore which is about 50 nm (Dimitrova et al., 2016; Gonen et al., 2012). Ndc80Cs were attached to the nanobead with the Ndc80C MT-binding domains oriented outward from the bead surface – Ndc80Cs *in situ* also take this orientation on the inner kinetochore and their distal ends directly bind MTs (Cheeseman et al., 2006; Ciferri et al., 2008; Wei et al., 2007). The Ndc80C was expressed in, and purified from, insect cells and visualized by its Spc24 component fused with GFP (Ndc80C-GFP) (Figure 1B).

In addition, Dam1C and Dam1C-GFP (in which the Dad1 component was fused with GFP) were expressed in, and purified from, bacterial cells (Figure 1B). Ndc80C-GFP was attached to a streptavidin-coated nanobead through a biotinylated anti-GFP nanobody (Ndc80C– nanobead; Figure 1C, top). Based on GFP intensity, we estimated that four Ndc80C-GFP molecules, on average, were attached to a single nanobead (Figure S1A). This was a slightly fewer than the 5-6 Ndc80Cs reportedly assembled on the MIND complexes at a single kinetochore (Dimitrova et al., 2016; Gonen et al., 2012). Dynamic MTs were generated *in vitro* from GMPCPP-stabilized MT seeds immobilized on coverslips, and observed by TIRF microscopy (Figure 1C, bottom). The Dam1C-GFP was able to track the end of depolymerizing MTs and accumulate there (up to 10–30 fold) (Figure S1B), as reported previously (Asbury et al., 2006; Tanaka et al., 2007; Westermann et al., 2006).

We investigated how Ndc80C (with GFP)–nanobeads behave with dynamic MTs and Dam1C (without GFP) *in vitro*. An Ndc80C–nanobead first attached to the lateral side of a MT (lateral attachment) (Figure 1D). When the laterally-attached MT depolymerized and its plus end caught up with the Ndc80C–nanobead, the nanobead became tethered at the MT plus end and subsequently tracked this MT end as it continued to depolymerize (end-on attachment) (Figure 1D). Nanobeads without Ndc80C did not bind the dynamic MTs themselves (Figure S1C). Thus, the Ndc80C–nanobeads on dynamic MTs recapitulated *in vitro* the behaviour of an authentic kinetochore in conversion from the lateral to end-on attachment on a single MT *in vivo* (Tanaka et al., 2007; Tanaka et al., 2005). In the presence of Dam1C, the end-on attachment continued in 96% cases (Figure 1D, F). In the absence of Dam1C, the end-on attachment could still be formed, but the Ndc80C–nanobead often (30% of cases) detached from the MT plus end following transient end-on interaction (Figure 1E, F), suggesting that Dam1C stabilizes end-on attachment. This is consistent with the important roles of Dam1C in interactions with Ndc80C *in vitro* (Lampert et al., 2010; Lampert et al., 2013; Sarangapani et al., 2013; Tien et al., 2010; Volkov et al., 2013) and in formation of end-on attachment of an authentic kinetochore to a MT *in vivo* (Kalantzaki et al., 2015; Maure et al., 2011; Tanaka et al., 2007).

### Direct comparison between end-on and lateral attachments suggests that the Dam1 C-terminus phosphorylation by Aurora B alters their relative strengths

The kinetochore–MT error correction relies on differential regulation of kinetochore interaction with the MT lateral side and with the MT end (Kalantzaki et al., 2015) (Figure 1A). However, there has been no assay to directly compare the strengths of the lateral and end-on attachments. Moreover, it is unknown whether the major outer kinetochore components Ndc80C and Dam1C are sufficient to explain such differential regulation. This prompted us to investigate how Ndc80C–nanobeads change their associated MTs *in vitro*. To this end, we observed situations where two MTs cross each other, one of which has an end-on attachment to an Ndc80C–nanobead during depolymerization (MT crossing assay; Figure 2A). This assay has two possible outcomes: the end-on attachment may continue and the Ndc80C– nanobead passes across the other MT (Figure 2A, blue rectangle); alternatively, the Ndc80– nanobead may be transferred from the end of the original MT to the lateral side of the other MT as the depolymerizing MT crosses it (Figure 2A, orange rectangle).

**Figure 2.**
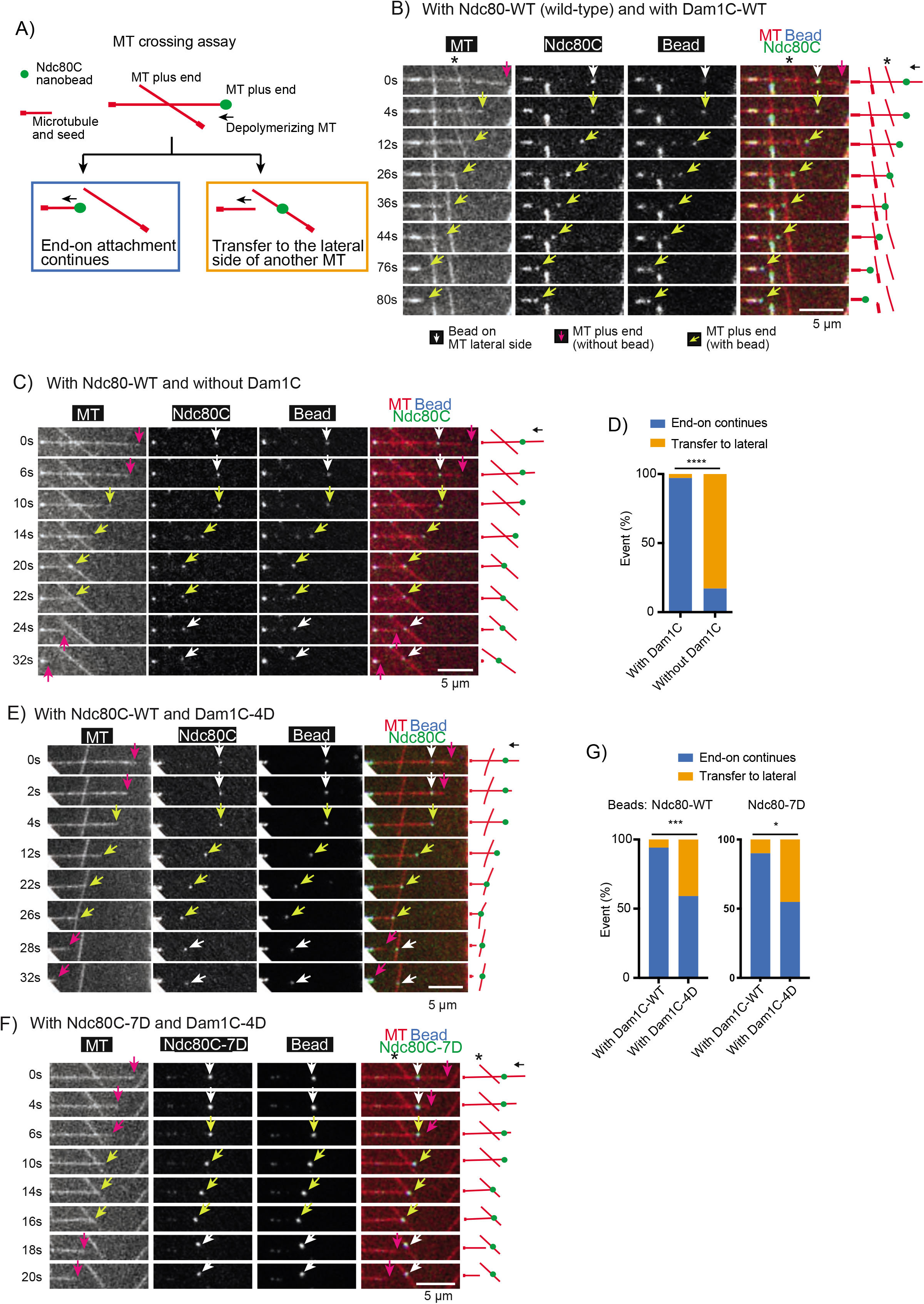
Direct comparison of strength between the MT lateral and end-on attachments to an Ndc80C-nanobead. **A**) Diagram explains the MT crossing assay. Two MTs cross each other, one of which forms end-on attachment to the Ndc80C–nanobead and depolymerizes. Two possible outcomes are shown; i) end-on attachment continues (blue rectangle) or ii) the Ndc80C–nanobead is transferred to the side of the other MT (orange rectangle). **B**) Images in time sequence show that the MT end-on attachment to the Ndc80C (wild-type)–nanobead continued after it had passed over the lateral side of another MT, in the presence of Dam1C (wild-type). Time 0 s was set arbitrarily. Scale bar, 5 μm. Also refer to diagrams (right). Asterisk indicates a crossing MT. **C**) Images in time sequence show that an Ndc80C (wild-type)–nanobead was transferred from the end of one MT to the side of another MT, in the absence of Dam1C. Time 0 s was set arbitrarily. Scale bar, 5 μm. Also refer to diagrams (right). **D**) Percentage of events in the MT crossing assay in the presence and absence of Dam1C (n=35 for each); i) continued end-on attachment of Ndc80C–nanobead (blue) and ii) transfer to the lateral side of another MT (orange). These events are shown in diagram A, inside the rectangles of the same colour. Difference between the two groups is significant (*p*<0.0001). **E, F**) Images in time sequence show that an Ndc80 (wild-type)–nanobead (E) or Ndc80C-7D– nanobead (F) was transferred from the end of one MT to the lateral side of another MT in the presence of Dam1C-4D. Scale bar, 5 μm. Also refer to diagrams (right). **G**) Percentage of events in the MT crossing assay with Ndc80 WT (wild-type)–nanobeads (left) or Ndc80-7D–nanobeads (right) with Dam1C-WT (wild-type) or Dam1C-4D (n=34, 44, 20 and 31 from left to right); i) continued end-on attachment (blue) and ii) transfer to the lateral side of another MT (orange). These events are shown in diagram A, inside the rectangles of the same colour. Difference between Dam1C-WT or Dam1C-4D is significant in the left and right graphs (left, *p*=0.0005; right, *p*=0.0125).

We conducted the MT crossing assay in the presence and absence of Dam1C (Figure 2B, C). In the presence of Dam1C, the end-on attachment continued in most cases (97%) when the Ndc80C–nanobead passed the other MT (Figure 2B, D). By contrast, in the absence of Dam1C, in most cases (83%) the Ndc80 nanobead was transferred to the side of the other MT (Figure 2C, D). Given that Dam1C accumulates at the end of depolymerizing MTs (Figure S1B), this difference can be explained by a weakened end-on attachment in the absence of Dam1C (Figure 1E). As a result, the affinity to the lateral attachment may surpass that of the end-on attachment, causing transfer of the Ndc80C–nanobead to the lateral side of another MT.

The Dam1C is the most important Aurora B substrate for error correction (Cheeseman et al., 2002; Kalantzaki et al., 2015) and its phosphorylation sites are clustered at the C-terminus of Dam1 protein, a component of the Dam1C. It is thought that error correction continues while Dam1 is phosphorylated and stops when bi-orientation is established and Dam1 is dephosphorylated (Keating et al., 2009). In addition to Dam1 phosphorylation, phosphorylation of the N-terminus of Ndc80 protein (a component of the Ndc80C) by Aurora B also contributes to error correction (Akiyoshi et al., 2009). To investigate how Dam1 and Ndc80 phosphorylation by Aurora B affects kinetochore–MT interaction *in vitro*, we expressed a) Dam1C carrying four phospho-mimic mutations at the C-terminus of Dam1 in bacteria and b) Ndc80C carrying seven phospho-mimic mutations at the N-terminus of Ndc80 in insect cells. The purified mutant Dam1Cs were called Dam1C-4D-GFP and Dam1C-4D, with and without GFP fusion to Dad1, respectively (Figure S2A). The mutant Ndc80Cs purified from insect cells were called Ndc80C-7D-GFP (with GFP fused to Spc24) (Figure S2A). Dam1C-4D-GFP tracked the plus end of a depolymerizing MT and accumulated there (Figure S2B, C) as did Dam1C (wild-type)-GFP (Figure S1B). Crucially, Dam1C-4D was able to support continuous end-on attachment of Ndc80C-WT (wild-type)–nanobeads in most cases without causing their detachment from the MT ends, as was Dam1C wild-type (Figure S2D). Meanwhile, when Ndc80C-7D-GFP were attached to a nanobead (Ndc80C-7D–nanobead), the nanobead showed lateral and end-on attachments to dynamic MTs, similarly to Ndc80C-WT–nanobeads (Figure 1D).

Subsequently we used Dam1C-4D and Ndc80-7D–nanobeads in the MT crossing assay (Figure 2A). In the presence of Dam1C-WT, the Ndc80C-WT and Ndc80C-7D –nanobead continued end-on attachment to a shrinking MT after crossing another MT in 94% and 90% cases respectively (Figure 2B, G). Whereas, in the presence of Dam1C-4D, Ndc80C-WT and Ndc80C-7D –nanobeads were directly transferred from the plus end of one MT to the lateral side of another MT in 41% and 45% cases, respectively (Figure 2E-G). We measured the angle between the two MTs, when an Ndc80C-WT–nanobead was transferred from the end of one MT to the lateral side of another in the presence of Dam1C-4D. Transfer occurred when two MTs crossed at a wide variety of angles ranging from 27° to 152° (Figure S2E). Similarly, Ndc80-7D–nanobead transfer occurred at a wide variety of angles.

These results suggest that the Dam1 phosphorylation by Aurora B kinase plays a key role in changing the relative strengths of the end-on and lateral attachments, i.e. end-on attachment is stronger in the absence of Dam1 phosphorylation, but it often becomes weaker than lateral attachment when Dam1 is phosphorylated. Our data suggest that Ndc80 phosphorylation may not make a major contribution to this process, which is consistent with the relatively minor roles of Ndc80 phosphorylation in error correction in yeast cells (Akiyoshi et al., 2009; Kalantzaki et al., 2015). Together with our previous observation in yeast cells (Kalantzaki et al., 2015), we suggest that the end-on attachment is specifically weakened by Dam1 phosphorylation by Aurora B, while the lateral attachment strength is unchanged, resulting in alteration in relative strengths of end-on and lateral attachments. This alteration likely drives the exchange of kinetochore–MT interactions, i.e. from end-on attachment on one MT to the lateral attachment on another MT, during error correction. Our *in vitro* system includes only the Ndc80C and Dam1C components of the kinetochore, thus showing that these two components sufficiently account for the differentially regulated end-on and lateral attachments during error correction.

### Evidence that phosphorylation of the Dam1 C-terminus by Aurora B kinase disrupts interaction between Dam1C and Ndc80C during error correction

How could the Dam1 C-terminus phospho-mimic mutants promote transfer of an Ndc80C– nanobead from the end of one MT to the lateral side of another in the MT crossing assay (Figure 2E–G)? The Dam1 C-terminus physically interacts with Ndc80C and its phosphorylation (or phospho-mimic mutants) weakens this interaction (Kalantzaki et al., 2015; Kim et al., 2017; Lampert et al., 2010; Sarangapani et al., 2013; Tien et al., 2010). However, it has been reported that the Dam1 C-terminus also interacts with MTs, and that its phosphorylation (or phospho-mimic mutants) weakens this interaction too (Legal et al., 2016; Zelter et al., 2015). Therefore, phospho-mimic mutants at the Dam1 C-terminus could disrupt either Dam1C–Ndc80C interaction or Dam1C–MT interaction when an Ndc80C–nanobead is detached from the MT end and is transferred to the lateral side of another MT.

To identify the point of disruption, we attached His-tagged Ndc80Cs (wild-type, without GFP; Figure 3A) to nanobeads (which are visible with fluorescence) using a biotinylated anti-His antibody (Figures 1C) and compared the behaviour of Dam1C-GFP and Dam1C-4D-GFP (Figures 1B, S2A) in the MT crossing assay (Figure 3B). The end-on attachment continued in most cases with Dam1C (wild-type)-GFP (Figure 3B, C), as with Dam1C (wild-type, without GFP) (Figure 2B, D). Moreover, Ndc80C nanobeads were often transferred to the lateral side of another MT with Dam1C-4D-GFP (Figure 3B, D), as with Dam1C-4D (without GFP) (Figure 2E–G)., We observed the location of Dam1C-4D-GFP shortly after this transfer occurred (Figure 3B, E). If the Dam1C–Ndc80C interaction were disrupted by Dam1 phospho-mimic mutants, Dam1C-4D-GFP signals would continue to track the end of a depolymerizing MT, thus moving away from the Ndc80C–nanobead that has been transferred to the side of another MT (Figure 3E, left). Alternatively, if the Dam1–MT interaction were disrupted by Dam1 phospho-mimic mutants, the Dam1C-4D-GFP would remain on the Ndc80C–nanobead after it is transferred to the lateral side of another MT (Figure 3E, right).

**Figure 3.**
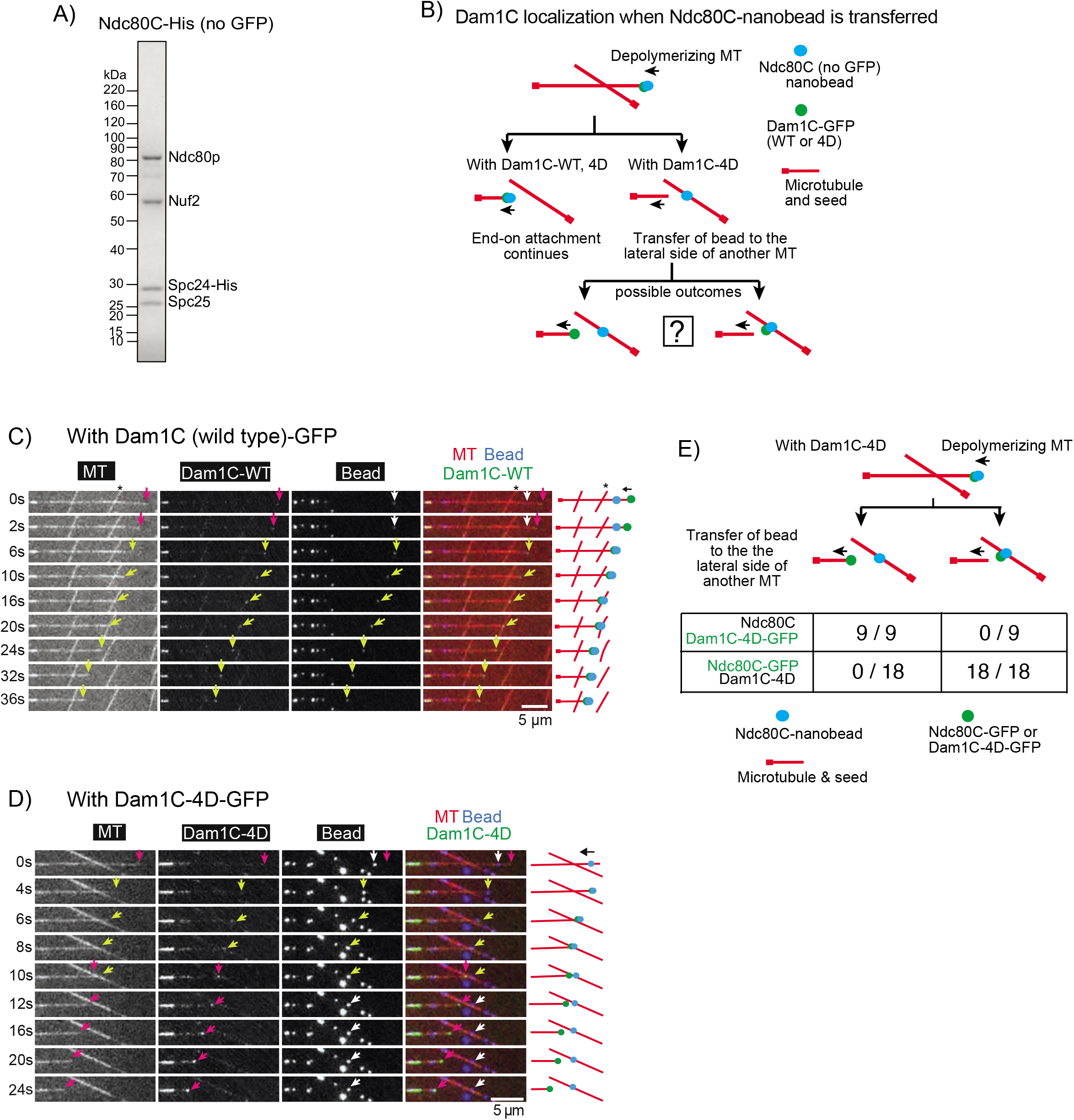
Dam1C-4D disrupts the interaction with Ndc80C, during the exchange of MTs attached to an Ndc80C–nanobead. **A**) Coomassie Blue stained gel showing purified Ndc80C-His (without GFP and with His-tag at the C-terminus of Spc24). **B**) Diagrams explains various outcomes in the MT crossing assay regarding the position of an Ndc80C–nanobead and the location of Dam1C-GFP (wild-type or 4D) signals. **C**) Images in time sequence show that the MT end-on attachment to an Ndc80C (wildtype)–nanobead continued after it had passed across the side of another MT in the presence of Dam1C (wild-type)-GFP. Dam1C (wild-type)-GFP signals were at the end of the depolymerizing MT with nanobead throughout the end-on attachment. Time 0 s was set arbitrarily. Scale bar, 5 μm. Also refer to diagrams (right). Asterisk indicates a crossing MT. **D**) Images in time sequence show that an Ndc80C (wildtype)–nanobead was transferred from the end of one MT to the side of another MT in the presence of Dam1C-4D-GFP. Dam1C-4D-GFP signals tracked the end of depolymerizing MT, moving away from the Ndc80C– nanobead. Time 0 s was set arbitrarily. Scale bar, 5 μm. Also refer to diagrams (right). **E**) Frequency of events in the MT crossing assay. The cases where Ndc80C–nanobeads were transferred to the lateral side of another MT, with Ndc80C (wildtype, no GFP) and Dam1-4D-GFP or with Ndc80C (wildtype)-GFP and Dam1C-4D (no GFP), were investigated. After transfer of the nanobead to the lateral side of another MT, Dam1-4D-GFP always tracked the end of the original MT (9 out of 9 events) while Ndc80C-GFP always located at the nanobead (18 out of 18 events). Difference between Dam1-4D-GFP and Ndc80C (wild-type)-GFP is significant (*p*<0.0001).

In all the cases where Ndc80C–nanobeads were transferred to the side of another MT, Dam1C-4D-GFP signals continued tracking the end of a depolymerizing MT, moving away from the Ndc80C–nanobead (Figure 3D, E). We could not detect any Dam1C-4D-GFP signals on Ndc80C-nanobeads after their transfer to the MT lateral side. For comparison, we looked at the location of Ndc80C-GFP signals soon after transfer of a nanobead to the side of another MT in the presence of Dam1C-4D (without GFP) (Figure 2E); in all these cases, Ndc80C-GFP signals remained on nanobeads after this transfer (Figure 3E). Thus, it can be ruled out that Dam1-4D proteins were present on nanobeads in a comparable amount of Ndc80C after the nanobead transfer. We conclude that phospho-mimic mutants at the Dam1 C-terminus disrupt the Dam1C–Ndc80C interaction, rather than the Dam1C–MT interaction, to promote transfer of an Ndc80C–nanobead from the end of one MT to the lateral side of another. This suggests that phosphorylation of the Dam1 C-terminus by Aurora B kinase disrupts the end-on attachment specifically by weakening interaction between Dam1C and Ndc80C during kinetochore–MT error correction.

### Kinetochore particles do not show direct transfer between two MTs in the presence of Dam1 and Ndc80 phospho-mimic mutants

The Ndc80C-nanobead system enabled direct comparison between the strengths of the endon and lateral attachments. Intriguingly, when Ndc80C–nanobeads were transferred from the end of one MT to the lateral side of another (with Dam1-4D phospho-mimic mutants), the end-on attachment was lost after the lateral attachment was formed, i.e. the Ndc80C– nanobeads were always attached to one or two MTs during the transfer (Figure 2E, F). In other words, they were ‘directly’ transferred between MTs. Such direct transfer possibly reflects the behaviour of native kinetochores during error correction. Indeed, Nicklas and colleagues implied that erroneous MT attachments are not released from the kinetochore until a new attachment is formed in grasshopper spermatocytes (Nicklas, 1997; Nicklas and Ward, 1994).

To address how native yeast kinetochores are transferred between MTs, we purified native kinetochore particles (KCp) from budding yeast, using a Flag tag fused to Dsn1 (a component of the kinetochore MIND complex) for immunoprecipitation (Akiyoshi et al., 2010). The KCp contained either wildtype Ndc80 (Ndc80-WT) or phospho-mimic mutant Ndc80-7D (at seven Aurora B phosphorylation sites in the N-terminus of Ndc80) with three copies of GFP at the Ndc80 C-terminus. Before purification of the KCp, we depleted Dam1 protein in yeast cells, in which an auxin-induced degron tag (aid) (Nishimura et al., 2009) was fused to the C-terminus of Dam1, by treatment with auxin (NAA). The western blots of the total cell lysates confirmed depletion of the most Dam1 protein after the NAA treatment (Figure 4A-i). As previously reported (Akiyoshi et al., 2010), the purified KCp contained a wide range of kinetochore components, including both inner and outer kinetochore components, but not Aurora B kinase (Figure 4A-ii, Table S1).

**Figure 4.**
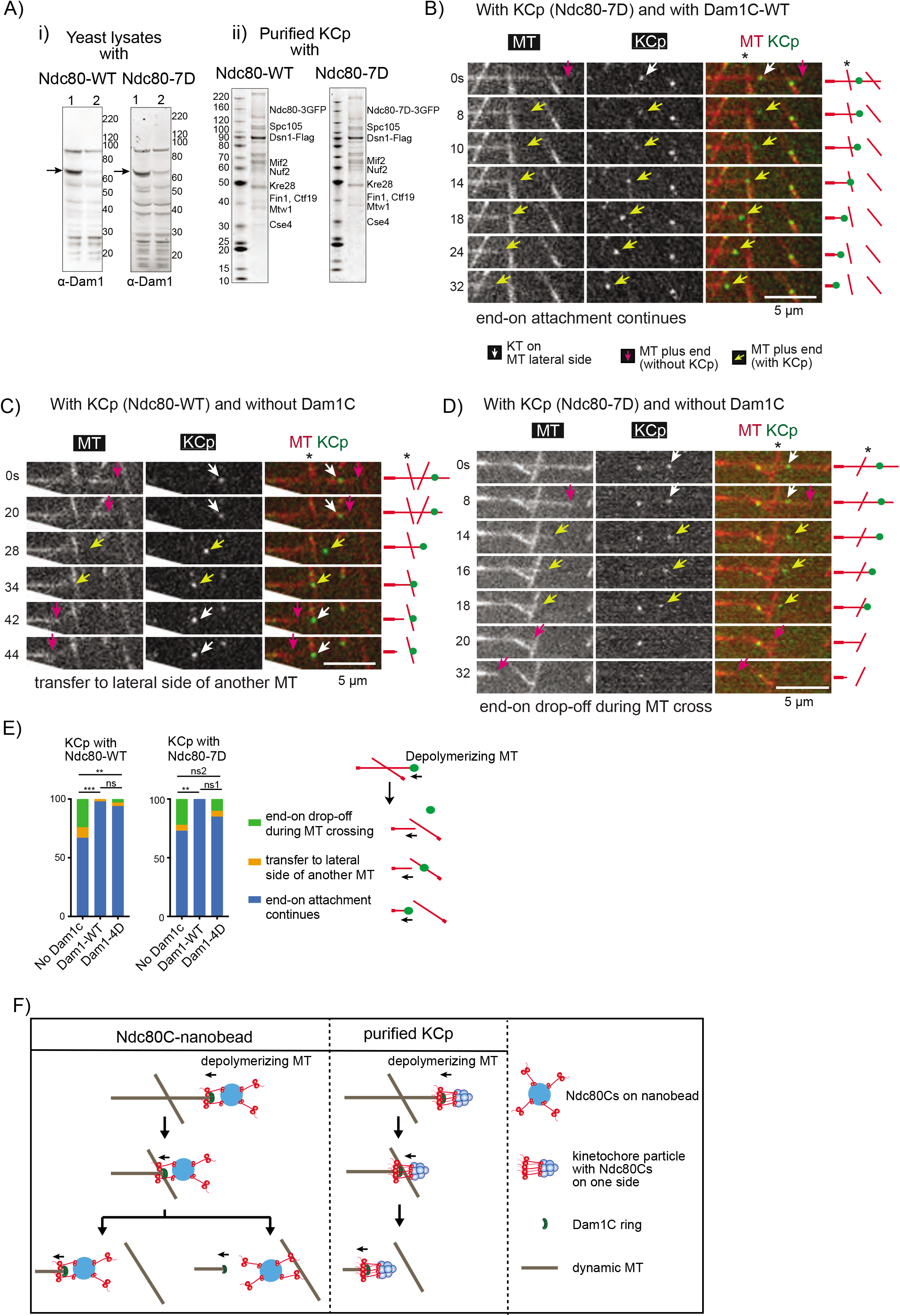
Behaviours of purified kinetochore particles (KCp) on dynamic MTs *in vitro*. **A**) (i) Western blots of yeast cell lysates from *dam1-aid* strains with *NDC80*-wild-type (WT) (left) and -7D (right). Cells were harvested before (lane 1) and after (lane 2) NAA treatment. The blots were probed with anti-Dam1p antibody. Arrows indicate Dam1-aid protein. Right: protein size markers (kDa). (ii) Purified KCp from *dam1-aid* strains with *NDC80* -WT (left) and -7D (right). Proteins were separated by SDS-PAGE and stained with SyproRuby. Proteins were identified by mass-spectrometry (Table S1) and are shown at predicted sizes. Left: protein size markers (kDa). **B**) Images in time sequence in a MT crossing assay shows that the MT end-on attachment to the KCp (purified from Dam1-depleted cells) continued after it had passed over the lateral side of another MT, in the presence of recombinant Dam1C. Also refer to diagrams (right). Asterisk indicates a crossing MT. Scale bar 5μm. In this example, KCp contained Ndc80-7D and recombinant Dam1C-WT was added to system. **C**) Images in time sequence show that the KCp (with Ndc80-WT, purified from Dam1-depleted cells) was transferred from the end of one MT to the lateral side of another MT (around 34 s), in the absence of recombinant Dam1C. **D**) Images in time sequence show that the KCp (with Ndc80-7D, purified from Dam1-depleted cells) detached from the end of a shrinking MT while passing over the lateral side of another MT (between 18-20s) in the absence of recombinant Dam1C. After the KCp had detached from the MT end, it was not visible by TIRF microscopy because it was no longer close to the coverslip. Keys are the same as in B). **E**) Percentage of various outcomes in the MT crossing assay for the KCp with Ndc80-WT (left) or Ndc80-7D (right). The KCp was purified from Dam1-depleted cells. The MT crossing assay was conducted in the absence of recombinant Dam1C or in the presence of recombinant Dam1C-WT or Dam1C-4D (from left to right n = 45, 44, 36 for the KCp with Ndc80-WT and n = 37, 30, 39 for the KCp with Ndc80-7D). Outcomes include i) continued end-on attachment (blue), ii) transfer to the lateral side of another MT (orange), iii) end-on drop-off during crossing of another MT (green). Also refer to the diagram (right). Comparisons between two Dam1 conditions give a) p=0.0005 (***), 0.53 (ns, not significant), 0.0086 (**) with Ndc80-WT and b) p= 0.0085 (**), 0.080 (ns1), 0.39 (ns2) with Ndc80-7D. **F**) Diagram shows the behaviours of Ndc80C-nanobeads and the purified KCp in the MT crossing assay. The Ndc80C-nanobead was often transferred to the lateral side of another MT in the presence of Dam1C-4D. By contrast, the most KCp continued to track the end of a shrinking MT while passing over the other MT, in the presence of Dam1C-4D. This difference may be explained by the difference in distribution and orientation of Ndc80Cs: The Ndc80Cs are randomly distributed around the 100 nm nanobeads and orient in all directions, whereas Ndc80Cs on the KCp may have a smaller footprint and orient mostly in one direction (Dimitrova et al., 2016; Gonen et al., 2012) (towards a MT).

We reconstituted KCp–MT interactions *in vitro* by mixing the purified KCp and recombinant Dam1C with dynamic MTs generated *in vitro*. The KCp first attached to the lateral side (lateral attachment) of dynamic MTs, then attached to the MT end, and tracked the end of shrinking MTs (end-on attachment) (Figure S3A). The continued end-on attachment relied on the Dam1C since its absence led to frequent (42–47%) detachment of KCp (containing either Ndc80-WT or -7D) from the end of shrinking MTs (end-on drop-off) (Figure S3B, C). With Dam1C-WT or Dam1C-4D, end-on drop-off of the KCp (with either Ndc80-WT or -7D) was rarely found (Figure S3C). Thus, the phospho-mimic mutants of Dam1 (Dam1-4D) and Ndc80 (Ndc80-7D in KCp) did not significantly affect continuous end-on attachment of the KCp; this is similar to their effects in the Ndc80C–nanobeads system (Figure S2D).

Next, we conducted the MT crossing assay with the KCp, as with Ndc80C–nanobeads. The KCp contained Ndc80-WT or 7D, and recombinant Dam1C was either present or absent in the system – where present, it contained either Dam1-WT or Dam1-4D. The assay gave three different outcomes as follows: 1) end-on attachment of KCp continued and passed a crossing MT (Figure 4B); 2) the KCp was transferred from the end of the original MT to the lateral side of another MT as the depolymerizing MT crossed it (Figure 4C); 3) the KCp detached from the MT end but was not transferred to another MT when the KCp reached it (end-on drop-off) (Figure 4D). Figure 4E shows frequency of these outcomes, in blue, orange and green bars, respectively.

In MT crossing assay with KCp (with Ndc80-WT), the absence of Dam1C led to more frequent transfer to the lateral side of another MT (Figure 4C; E, orange, 9%) and end-on drop-offs (Figure 4E, green, 24%) in comparison with the presence of Dam1C-WT. Also, KCp with Ndc80-7D showed similar outcomes in the absence of Dam1C (Figure 4D, E). In the presence of recombinant Dam1C-WT and -4D, end-on attachment continued in most cases (Figure 4B, E) and the other two events were rare. The outcomes were similar irrespective of Ndc80-WT or -7D in the KCp (Figure 4E). In summary, phospho-mimic mutants of either Dam1 or Ndc80 showed no significant effects on the behaviour of the KCp in the MT crossing assay. With Dam1C-4D, on rare occasions, the KCp was transferred from the end of one MT to the lateral side of another, in contrast to the behaviour of Ndc80C-nanobeads.

We suspect that this difference in behaviour between the Ndc80C–nanobeads and the KCp is due to different distributions and orientations of the Ndc80Cs, i.e. Ndc80Cs are randomly distributed around the 100 nm nanobeads and orient in all directions, whereas Ndc80Cs on the KCp may have a smaller footprint and orient mostly in one direction (towards a MT) (Dimitrova et al., 2016; Gonen et al., 2012) (Figure 4F). The results with purified KCp suggest that direct transfer between MTs, without generating completely unattached kinetochores, may not be a feature of authentic kinetochores during error correction, at least, in budding yeast.

## Discussion

During the process of establishing chromosome bi-orientation, sometimes aberrant interactions are formed between kinetochores and MTs. In such cases, these interactions must be detected, dissolved and re-formed in a process called error correction, in which Aurora B kinase plays central roles (Hauf et al., 2003; Lampson et al., 2004; Tanaka et al., 2002). Measurement of the end-on attachment of kinetochores to MTs using optical tweezers has demonstrated that end-on attachment is weakened by the action of Aurora B (Sarangapani et al., 2013). We have shown that, while Aurora B weakens the end-on attachment, the lateral attachment is impervious to Aurora B regulation, i.e. the end-on and lateral attachments are differentially regulated (Kalantzaki et al., 2015). This led us to propose the model that, during error correction, an end-on attachment is disrupted by the action of Aurora B and then replaced by lateral attachment to a different MT, which is then converted to end-on attachment (Kalantzaki et al., 2015) (Figure 1A).

However, for this model to work, Aurora B needs to change the relative strengths of the endon and lateral attachments. Here, we directly compared the strength of end-on and lateral attachments by reconstituting a kinetochore–MT interface *in vitro* using Ndc80C–nanobeads and Dam1C-loaded dynamic MTs. Our results suggest that without Aurora B activity end-on attachment is stronger than lateral attachment, but this relative strength is reversed when Dam1C is phosphorylated by Aurora B (Figure 2).

Previous works have shown that Dam1C and Ndc80C are important targets of Aurora B in promoting error correction in budding yeast (Akiyoshi et al., 2009; Cheeseman et al., 2002). However, it has been unclear whether the kinetochore–MT interface formed by Ndc80C and Dam1C is sufficient to account for differential regulation of end-on and lateral attachments by Aurora B kinase or whether any additional kinetochore components are involved. The Ndc80C–nanobead assay uses only recombinant Ndc80C and Dam1C components of the kinetochore, demonstrating that these two components are sufficient for the differential regulation of end-on and lateral attachments by Aurora B (Figure 2).

Previous results have also suggested that Dam1 C-terminus phosphorylation by Aurora B weakens Dam1C–Ndc80C interactions (Kalantzaki et al., 2015; Kim et al., 2017; Lampert et al., 2010; Sarangapani et al., 2013; Tien et al., 2010) as well as Dam1C–MT interactions (Zelter et al., 2015). However, it is disputed which is most critical when end-on attachment is disrupted by Aurora B. By visualizing Dam1C in the Ndc80C–nanobead assay, we showed that Dam1C–Ndc80C is the major disruption point when end-on attachment is lost by Dam1 C-terminus phosphorylation by Aurora B (Figure 3).

When Ndc80C–nanobeads were transferred from the MT end to the lateral side of another MT in the presence of Dam1 phospho-mimic mutants, this transfer was direct, i.e. the end-on attachment was lost only after the lateral attachment was formed. Does this reflect the behaviour of authentic kinetochores during error correction? In fact, it was previously implied that erroneous MT attachments are not released from the kinetochore until a new attachment is formed in grasshopper spermatocytes (Nicklas, 1997; Nicklas and Ward, 1994). However purified kinetochore particles from yeast cells rarely showed such a direct transfer between MTs, in contrast to the Ndc80C–nanobeads (Figure 4), suggesting that it may not be a behaviour of authentic kinetochores, at least in budding yeast. We suspect that the difference between the Ndc80C–nanobeads and the kinetochore particles is due to difference in distribution and orientation of the Ndc80Cs (Figure 4F).

If error correction of kinetochore–MT interactions does not involve direct transfer of kinetochores between MTs, disruption of end-on attachment may occur before a lateral attachment to another MT is formed. However, Dam1 phospho-mimic mutants rarely showed detachment of either Ndc80–nanobeads or purified kinetochore particles from a MT end without a crossing MT, even if the end-on attachment was weakened (Figure S2D, S3C). Moreover, Dam1 phospho-mimic mutants showed only slow (over 30 min or longer) kinetochore detachment from the MT end in cells (Kalantzaki et al., 2015). To explain these observations, we speculate that rapid disruption of the end-on attachment may happen only in the context of aberrant kinetochore–MT interactions such as syntelic attachment where both sister kinetochores interact with MTs from the same spindle pole. For example, if two MTs, attached to sister kinetochores, have different dynamics, the resulting twisting force would disrupt the end-on attachment of one of sister kinetochores (Ault and Rieder, 1992). Such a mechanism would prevent both sister kinetochores simultaneously losing MT attachment and therefore avoid a chromosome drifting away from the spindle during error correction.

In conclusion, our study suggests that Ndc80C and Dam1C are sufficient to constitute key regulation of kinetochore–MT interactions by Aurora B kinase. Dam1 phosphorylation by Aurora B weakens the association with the Ndc80 complex, disrupts end-on attachment and promotes the exchange to a new MT lateral attachment. Such exchange of kinetochore–MT interactions promotes error correction to establish bi-orientation.

## Acknowledgements

We thank Tanaka lab members, E.R. Griffis, E.A. Katrukha, R. Cross, D. Peet, T. Surrey, G. Ball, M. Gierlinski and M. Miller for discussion; S. Biggins for the kinetochore particle purification protocol; S. Harrison, T. Davis, T. Richmond, K. Nakayama, L. Garcia and Addgene for reagents; E.R. Griffis, P. Appleton, S. Swift, D. Bajer and Dundee Imaging Facility for TIRF microscope setting and maintenance; Dundee FingerPrints Proteomic Facility for mass spectrometry analyses; N. Sheidaei for help in protein purification; and the OMERO team for technical help. This work was supported by the Wellcome Trust (096535/Z/11/Z, 097945/B/11/Z, 219418/Z/19/Z) and Medical Research Council (K015869). The authors declare no competing financial interests.

## Methods

### Plasmid constructs

To express and purify *S. cerevisiae* Ndc80C-GFP, *NDC80, GST-NUF2, SPC24-GFP* and *SPC25* were cloned into Multibac vectors, pFL (*NDC80* and *GST-NUF2*) and pUCDM (*SPC24-GFP* and *SPC25*) (Bieniossek et al., 2008). A PreScission cleavage site was inserted between *GST* and *NUF2*. The two plasmids (pT3044 and pT3045, respectively) were then fused by Cre-lox recombination *in vitro*, making pT3046. After transfection of pT3046 into DH10MultiBac *E. coli* (Bieniossek et al., 2008) bacmid DNA was prepared using the alkaline lysis method. To express and purify Ndc80C-7D-GFP, *NDC80* in pT3044 was replaced with *NDC80-7D* mutant, in which Thr 21, Ser 37, Thr 54, Thr 71, Thr 74, Ser 95 and Ser 100 of the *NDC80* N-terminal region were replaced with aspartates (Akiyoshi et al., 2009; Kalantzaki et al., 2015), making pT3352. pT3352 and pT3045 were fused in vitro to make pT3371 expressing Ndc80C-7D-GFP. To express and purify Ndc80C (without GFP), pT3045 was modified by replacing GFP on *SPC24-GFP* with a His tag (*SPC24-His*), making pT3219. pT3044 and pT3219 were fused, making pT3220 with Ndc80C without GFP. The MultiBac system was a gift from Prof. Tim Richmond.

To express *S. cerevisiae* Dam1C and Dam1C-GFP components in *E. coli*, plasmid constructs were obtained from Prof. Stephen Harrison and Prof. Trisha Davis (Gestaut et al., 2008; Miranda et al., 2005), respectively. These constructs express Spc34 tagged with His at its C-terminus. To express Dam1C-4D and Dam1C-4D-GFP components, plasmid constructs were generated by DNA synthesis (DC Biosciences, Dundee) and cloning to replace four serine residues at C-terminus of Dam1 (serines at 257, 265, 292, 327 positions of Dam1) with aspartates (Cheeseman et al., 2002; Kalantzaki et al., 2015) (pT3143 and pT3145, respectively).

### Protein purification

Bacmid DNA with Ndc80C components was transfected into Sf9 insect cells to produce baculovirus. For protein expression, Sf9 cells were grown for 60 to 72 hrs at 27 °C. The cells were then harvested and washed with PBS (8 mM Na2HPO4, 2mM KH2PO4, 2.7mM KCl, 137mM NaCl, pH7.4) and stored at −80 °C. For purification of Ndc80C proteins, the cells were resuspended in buffer containing 50 mM Hepes pH 7.4, 300 mM NaCl, 1 mM EDTA, 10% Glycerol, 1% Triton X-100, 1 mM DTT and protease inhibitor cocktail (Roche), and lysed with a Dounce homogenizer and sonication at 4 °C. The cell lysate was clarified by centrifugation at 25 000 x g for 45 to 60 min. The soluble fraction was bound to the GST Sepharose 4B (GE Healthcare) for 90 to 120 min at 4 °C. The unbound fractions were removed on a gravity flow column and washed with buffer containing 50 mM Hepes pH 7.4, 250 mM NaCl, 1 mM EDTA, 10% Glycerol and 1 mM DTT. The proteins were eluted in buffer containing 50 mM Hepes pH 7.4, 150 mM NaCl, 1 mM EDTA, 0.05% Triton X-100 and 1 mM DTT, by cleavage with PreScission protease (GE Healthcare) which cleaves between GST and Nuf2 in the GST-Nuf2 fusion protein. The eluted proteins were further purified by gel filtration using Superose 6 Increase 10/300 GL column (GE Healthcare) equilibrated with buffer containing 25 mM Tris-Cl pH 7.6, 250 mM NaCl, 1 mM EDTA, 3 mM MgCl_2_, 5% Glycerol and 1mM DTT. The fractions containing Ndc80C were pooled, and the buffer was exchanged into the one containing 80 mM Pipes pH 6.8, 1 mM MgCl_2_, 1 mM EGTA, 150 mM KCl, 5% sucrose and 0.2 mM DTT, using a PD-10 desalting column (GE Healthcare). Purified Ndc80C was stored at −80°C.

For purification of Dam1C proteins, Rosetta™ 2(DE3) cells (Novagen 71401) transformed with respective constructs were grown at 37 °C until OD_600_ reached 0.6–0.7. Then protein expression was induced by 0.1 mM IPTG for 22 hrs at 15-16 °C. The cells were harvested and stored at –80 °C. Dam1C proteins were purified in a three-step process, by affinity, ion-exchange and gel filtration chromatography as described (Miranda et al., 2005). The bacterial cells were resuspended in ice-cold buffer containing 50 mM sodium phosphate pH 7.5, 500 mM NaCl, 1 mM EDTA, 0.5% Trition X-100, 20 mM imidazole, 1 mM DTT and protease inhibitor cocktail (Roche). Cells were then lysed on ice and sonicated at 4°C, and the cell debris was separated by centrifugation at 25 000 x g for 45–60 min. The supernatant was incubated with Ni-NTA agarose (Qiagen) at 4 °C for 90 min. Unbound fractions were separated using a gravity flow column. Protein-bound Ni-NTA agarose was washed with buffer containing 50 mM sodium phosphate pH 7.5, 500 mM NaCl, 1 mM EDTA, 0.5% Trition X-100, 50 mM imidazole and 1 mM DTT. Then proteins were eluted with the same buffer containing 250 mM imidazole, without Triton X-100. The eluted proteins were exchanged into 50 mM sodium phosphate pH7.5, 100 mM NaCl, 1 mM EDTA, 0.2 mM DTT by using PD-10 desalting columns (GE Healthcare), then 1 mM ATP and 250-fold molar excess of synthetic peptide NRLLTG was added and incubated for 1 h at 4 °C. Proteins were purified on a MonoQ 5/50 column (GE Healthcare) with a gradient of 100 to 1000 mM NaCl. The eluent was mixed again with 1 mM ATP and NRLLTG peptide (250-fold molar excess) and incubated for 1 h at 4 °C and purified on a Superose 6 Increase 10/300 GL column (GE Healthcare) equilibrated with 25 mM sodium phosphate pH7.4, 500 mM NaCl, 1 mM EDTA and 0.2 mM DTT. The fractions corresponding to Dam1C proteins were pooled and the buffer was exchanged into the one containing 80 mM PIPES pH 6.8, 1 mM MgCl_2_, 1 mM EGTA, 150 mM KCl, 5% sucrose and 0.2 mM DTT. Purified Dam1C was stored at −80 °C.

### Attaching Ndc80C to nanobeads

Fluorescent (excitation at 647 nm) streptavidin-coated magnetic nanobeads (100 nm diameter; will be referred as nanobeads hereafter) were obtained from Creative Diagnostics, USA (cat No.WHM-ME647). Ndc80C-GFP, Ndc80C-7D-GFP and Ndc80C-His were attached to nanobeads using biotinylated anti-GFP nanobody (Chromotek cat No. gtb-250) and biotinylated anti-His antibody (Quiagen cat No. 34440), as appropriate, as shown in Figure 1C (top). For this, the beads were incubated with biotinylated anti-GFP nanobody (or biotinylated anti-His antibody) for about 1 h along with 5 mg/mL BSA at 4 °C. The unbound fractions were removed after the nanobeads were bound to the magnet and washed three times in MRB80 buffer (80 mM Pipes pH 6.8, 1 mM MgCl_2_, 1 mM EGTA) supplemented with 5 mg/ml BSA. The nanobeads were then incubated with Ndc80C in MRB80 buffer supplemented with 5 mg/ml BSA for 1 h at 4 °C, washed three times and finally resuspended in MRB80 buffer.

### Generation of dynamic MTs on coverslips

Purified tubulin proteins were obtained from Cytoskeleton, Inc. For preparation of MT seeds, 20 μM of porcine tubulin mix containing 18% biotinylated tubulin, 12% rhodamine-labelled tubulin and 70% unlabelled tubulin was incubated with 1 mM guanylyl-(α,β)-methylenediphosphonate (GMPCPP) on ice and subsequently at 37 °C for 30 min. MTs were separated from free tubulin by centrifugation using an Airfuge (Beckman Coulter) for 5 min. The MTs were subjected to another round of depolymerization and polymerization with 1 mM GMPCPP, and the final MT seed samples were stored in MRB80 buffer (80 mM Pipes pH 6.8, 1 mM MgCl_2_, 1 mM EGTA) supplemented with 10% glycerol.

Coverslips were plasma cleaned using Carbon Coater (Agar Scientific) and treated with PlusOne Repel-Silane (GE Healthcare) for 10–15 min. The coverslips were further cleaned by sonication in methanol and finally rinsed in water. Flow chambers were assembled with cleaned coverslips and microscopy slides using double-sided tape.

The chambers were treated with 0.2 mg/ml PLL-PEG-biotin (Surface solutions, Switzerland) in MRB80 buffer for 5 min. Subsequently they were washed with buffer and incubated with 1 mg/ml NeutrAvidin (Thermo Fisher) for 5 min. MT seeds were attached to the coverslips with the biotin-NeutrAvidin links and then incubated with NeutrAvidin once again to neutralize the exposed biotins on MT seeds that were already bound to coverslips. Finally, the chambers were incubated with 1 mg/ml κ-casein.

The *in vitro* reaction mixture was prepared in MRB80 buffer (80 mM Pipes pH 6.8, 1 mM MgCl_2_, 1 mM EGTA) consisting of 12 μM tubulin mix (11.5 μM unlabelled-tubulin and 0.5 μM rhodamine-tubulin), 50–60 mM KCl, 2 mM MgCl_2_, 0.1% Methylcellulose, 0.5 mg/ml κ-casein, 1 mM GTP, 6 mM DTT, oxygen scavenging system (400 μg/ml glucose-oxidase, 200 μg/ml catalase, 4 mM DTT and 20 mM glucose) and 10 nM of relevant Dam1C proteins. The mixture was centrifuged for 5 min using Airfuge. To the supernatant, nanobeads coated with Ndc80C-GFP, Ndc80C-7D-GFP or Ndc80C-His (or purified KCp) were mixed and added to the flow chamber containing MT seeds. The chamber was sealed with vacuum grease and observed at 30 °C by TIRF microscopy. To study behaviour of Dam1C on dynamic MTs, the reaction mixture was prepared in the same way as above but without nanobeads.

### TIRF Microscopy and Image analysis

Images of dynamic MTs were acquired by TIRF microscopy using Nikon Eclipse Ti-E (Nikon) inverted research microscope equipped with four diode lasers (405 nm, 488 nm, 561 nm, 647 nm; Coherent), AOTF shutter (Solamere Technology), appropriate filters (Chroma), perfect focus system, the CFI Apochromat TIRF 100X 1.49 N.A. oil objective lens (Nikon) and Evolve Detla EMCCD 512×512 camera (Photometrics). The TIRF system was controlled with μ-manager software (Open Imaging) (Edelstein et al., 2014). A temperature control chamber (Okolab) was used to maintain the temperature.

Images were analysed using ImageJ and OMERO. Kymographs were generated in time sequence along a chosen line for an individual MT, using KymoResliceWide plugin on ImageJ. The Dam1C-GFP intensity at the end of depolymerizing MT was obtained from Kymographs along the path of the MT depolymerisation, using imageJ. Statistical analyses were carried out with Prism software (GraphPad), using Fishers exact test or Chi-square test The fluorescence intensity of GFP protein and that of Ndc80C-GFP on nanobeads were obtained from TIRF microscopy images (in semi TIRF angle) and analyzed using ImageJ plugin DoM v.1.1.6 (Detection of Molecules, https://github.com/ekatrukha/DoM_Utrecht). All the experiments were repeated at least twice and similar results were obtained.

### Yeast strains

Two yeast strains (T12525 and T13763, see their genotypes below) were constructed for purification of kinetochore particles (see next section), as follows: Three copies of *FLAG* tags with *KAN* marker (pT2329) were added to the *DSN1* C-terminus, three copies of *GFPs* with KAN marker (Maekawa et al., 2003) were added to the *NDC80* C-terminus, and auxin-inducible degron (aid) tag with *clonNAT* marker (Nishimura et al., 2009) was added to the *DAM1* C-terminus, at their original loci. To do so, these tags and selection markers were amplified by PCR and yeast cells were transformed with the PCR products. The construct of rice *TIR1* under *ADH1* promoter was previously reported (Nishimura et al., 2009) and integrated at *TRP1* locus. These constructs were introduced into yeast cells by sequential transformation or gathered in a single yeast strain by crossing strains with each construct. To replace *NDC80* (wild-type) in T12525 with *NDC80-7D* and to make T13763, 5’ promoter DNA fragment of *NDC80, HIS* marker, 5’ promoter and *Ndc80-7D* ORF were cloned in this order to make the replacement cassette pT2037.

T12525: *MATa dsn1::DSN1-3XFLAG::KAN-MX4 ndc80::NDC80-3xGFP::KAN dam1::dam1-aid::NAT-NT2 trp1::ADH1p-TIR1-9Myc::TRP1*

T13763: *MATa dsn1::DSN1-3XFLAG::KAN-MX4 ndc80::HIS3::Ndc80-7D-3xGFP::KAN dam1::dam1-aid::NAT-NT2 trp1::ADH1p-TIR1-9Myc::TRP1*

### Purification of kinetochore particles

Kinetochore particles were affinity purified using FLAG-tag on Dsn1 as described in (Akiyoshi et al., 2010; Gupta et al., 2018) after depletion of Dam1 protein. Briefly, overnight grown yeast cells (T12525 or T13763) were diluted to 0.1 OD_600_ in YPAD (1% yeast extract, 2%peptone, 0.01% adenine hydrochloride, 2% glucose) medium and grown further at 25 °C. When the OD_600_ of the culture reached 0.6, cells were treated with 1mM NAA (1-Naphthaleneacetic acid) for 90 min to deplete Dam1p tagged with auxin-induced degron. The cells were harvested by centrifugation and washed with MilliQ water and 0.2mM PMSF (phenylmethylsulfonyl fluoride) and then washed with buffer H (25mM HEPES pH8.0, 150mM KCl, 2mM MgCl_2_, 0.1mM EDTA pH8.0, 0.5mM EGTA pH8.0, 0.1% NP-40, 15% glycerol) containing protease inhibitors and phosphatase inhibitors. The cells were resuspended in lysis buffer containing Buffer H supplemented with protease and phosphatase inhibitors, dropped into liquid nitrogen by pipetting to create “dots” and stored at −80 °C. The frozen “dots” were ground into powder in a freezer mill for 2 min and cooled down for 2 min – this cycle was repeated 7 times. The cell powder was thawed on ice and incubated with benzonase (300 units/ml) for 30 min at 4 °C. The soluble fraction was separated by centrifugation at 20000g for 30 min and the supernatant was centrifuged again at 72000 xg for 60 min. The final supernatant was incubated with FLAG antibody conjugated beads (anti-FLAG M2 magnetic beads #M8823 from Sigma) for 3 hrs with constant rotation. The beads were washed four times with buffer H containing protease inhibitors, phosphatase inhibitors and 2mM dithiothreitol (DTT) and washed three more times with buffer H containing protease inhibitors and phosphatase inhibitors. The bound proteins were eluted in buffer HE (25mM HEPES pH8.0, 150mM KCl, 2mM MgCl_2_, 0.1mM EDTA pH8.0, 0.5mM EGTA pH8.0, 15% glycerol) containing protease inhibitors, phosphatase inhibitors and 0.5 mg/ml FLAG peptides. The eluted proteins were aliquoted and stored at −80 °C.

### Western blots

Proteins were separated on an SDS-PAGE gel and transferred to a nitrocellulose membrane using iBlot 2 apparatus (ThermoFisher Scientific). The membrane was incubated in TBST (20mM Tris, 150mM NaCl, 0.1% Tween 20) containing 5 % milk for 1 h, washed three times with TBST and incubated with the primary antibody – the polyclonal anti-Dam1p (Keating et al., 2009) or monoclonal anti-GFP antibody (#11814460001 from Sigma) overnight at 4 °C. The blots were incubated with either HRP-tagged or IRDye^®^ secondary antibody. The blots were then scanned using either with ChemiDoc (Bio-Rad) or LI-COR odyssey imager.

### Mass spectrometry

Kinetochore particles were purified as described above and eluted in buffer containing 0.2% RapiGest SF surfactant (Waters^®^) and 50mM HEPES pH8.0. The protein sample (~150ng) was digested in solution with trypsin overnight at 30 °C, processed with HiPPR Detergent Removal Spin Column Kit (ThermoFisher Scientific #88305) and sample was dried by SpeedVac at room temperature and stored at −20 °C.

LC-MS analysis was done at the FingerPrints Proteomics Facility (University of Dundee). Analysis of peptide readout was performed on a Q Exactive™ plus, Mass Spectrometer (Thermo Scientific) coupled with a Dionex Ultimate 3000 RS (Thermo Scientific). LC buffers used were buffer A (0.1% formic acid in Milli-Q water) and buffer B (80% acetonitrile and 0.1% formic acid in Milli-Q water). Samples were resuspended in 35μl of 1% formic acid and 15 μl were loaded onto a trap column (PepMap nanoViper C18 column, 5 μm, 100 Å, Thermo Scientific) equilibrated in 0.1% TFA. The trap column was washed for 5 min at the same flow rate with 0.1% TFA and then switched in-line with a Thermo Scientific, resolving C18 column (PepMap RSLC C18 column, 2 μm, 100 Å). The peptides were eluted from the column at a constant flow rate of 300 nl/min with a linear gradient from 2% buffer B to 5 %, then to 35%, and finally to 98%. The column was then washed with 98% buffer B for 20 min and reequilibrated in 2% buffer B for 17 min. The column was kept at a constant temperature of 50°C. Q-exactive plus was operated in data dependent positive ionization mode. The source voltage was set to 2.5 kV and the capillary temperature was 250 °C. A scan cycle comprised MS1 scan (m/z range from 350-1600, ion injection time of 20 ms, resolution 70 000 and automatic gain control 1×10^6^) acquired in profile mode, followed by 15 sequential dependent MS2 scans (resolution 17500) of the most intense ions fulfilling predefined selection criteria. The HCD collision energy was set to 27% of the normalized collision energy. Mass accuracy was checked before the start of samples analysis. For protein identification, MaxQuant version 1.6.0.16 (Tyanova et al., 2015) was run against S. cerevisiae protein database.

---

**Figure S1.**
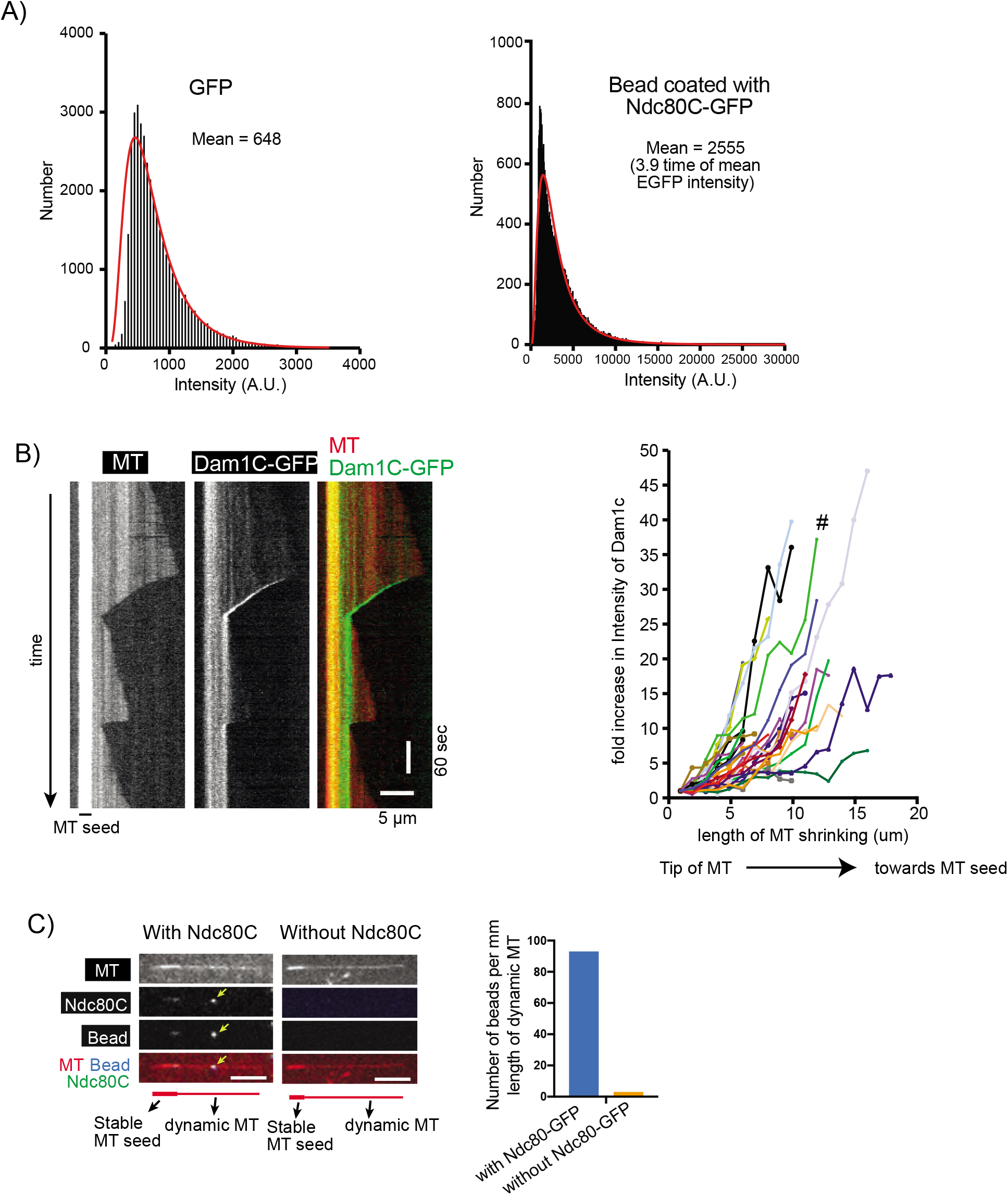
Supplemental data associated with Figure 1. A) Signal quantification of GFP (by itself, not fused to Ndc80C) and Ndc80C-GFP-coated beads. Black bars (bin size 50) represent the number of fluorescent spots with the indicated intensity. The red line is the lognormal fit of the quantification. A.U., arbitrary unit. B) Kymograph (left) shows that the Dam1C-GFP signal tracked the end of a depolymerizing MT. Scale bars, 5 μm (horizontal) and 60 s (vertical). Graph (right) shows fold increases of Dam1C-GFP signals at the plus ends of individual MTs. The Dam1C-GFP signal for the first 1 μm after their appearance was averaged and set to one for normalization. The Dam1C signal during subsequent MT shrinkage was averaged in 1-μm increments, normalized and plotted against the length of MT shrinkage. # shows the fold increase of the example shown in the kymograph (left). C) Images show a representative example of dynamic MTs when they were incubated with Ndc80C-GFP-coated nanobeads (left) or control nanobeads without Ndc80C-GFP (right). The yellow arrowhead indicates an Ndc80C-GFP-coated nanobead on the lateral side of a dynamic MT. Size bar, 5 μm. Graph shows the number of nanobeads (with and without Ndc80C-GFP coating) per milli meter length of dynamic MTs.

**Figure S2.**
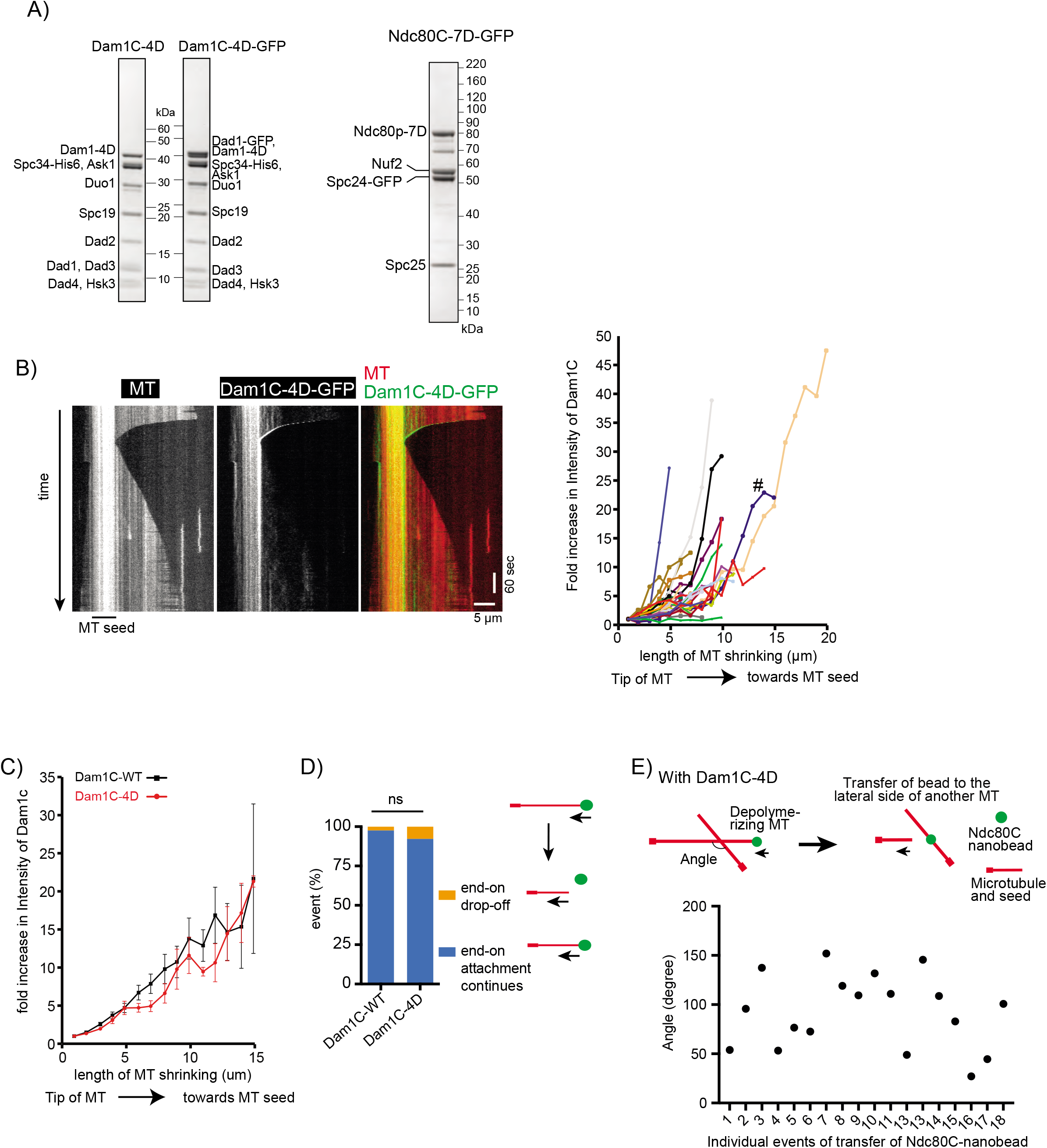
Supplemental data associated with Figure 2. A) Coomassie Blue stained gel (left panel) shows purified Dam1C-4D (in which four serine residues at the C-terminus of Dam1 were replaced with aspartate) with and without GFP at the C-terminus of Dad1. Coomassie Blue stained gel (right panel) shows purified Ndc80C-7D-GFP (with seven serine residues at N-terminus of Ndc80 replaced with aspartate). B) Kymograph (left) shows that Dam1C-GFP-4D signals tracked the end of a depolymerizing MT. Scale bar, 5 μm (horizontal) and 60 s (vertical). Graph (right) shows fold increases of Dam1C-GFP signals at the plus ends of individual MTs, which were obtained and plotted as in Figure S1B. # shows the fold increase in the example shown in the kymograph (left). C) Dam1C (wild-type)-GFP and Dam1C-4D-GFP show similar fold increases at the plus end of shrinking MTs. The fold increase of Dam1C (wild-type)-GFP (n=28) (black squares) or Dam1C-4D-GFP (n=32) (red circles) signals at the shrinking MT ends (Figure S1B and S2B) was averaged among multiple MTs and plotted against the length of MT shrinkage. Error bars show SEM. D) Graph shows the percentage of events; end-on drop-off and continuous end-on attachment, observed in the presence of Dam1C wild-type (WT) (n=122) and Dam1C-4D (n=90). Difference between the two groups is not significant (*p*=0.10). E) Angles made by two MTs between which Ndc80C (wild-type)–nanobeads were transferred, from end-on to the lateral side of another MT, in the presence of Dam1C-4D. Angles were measured in individual events as shown in diagram (top) and plotted (bottom).

**Figure S3.**
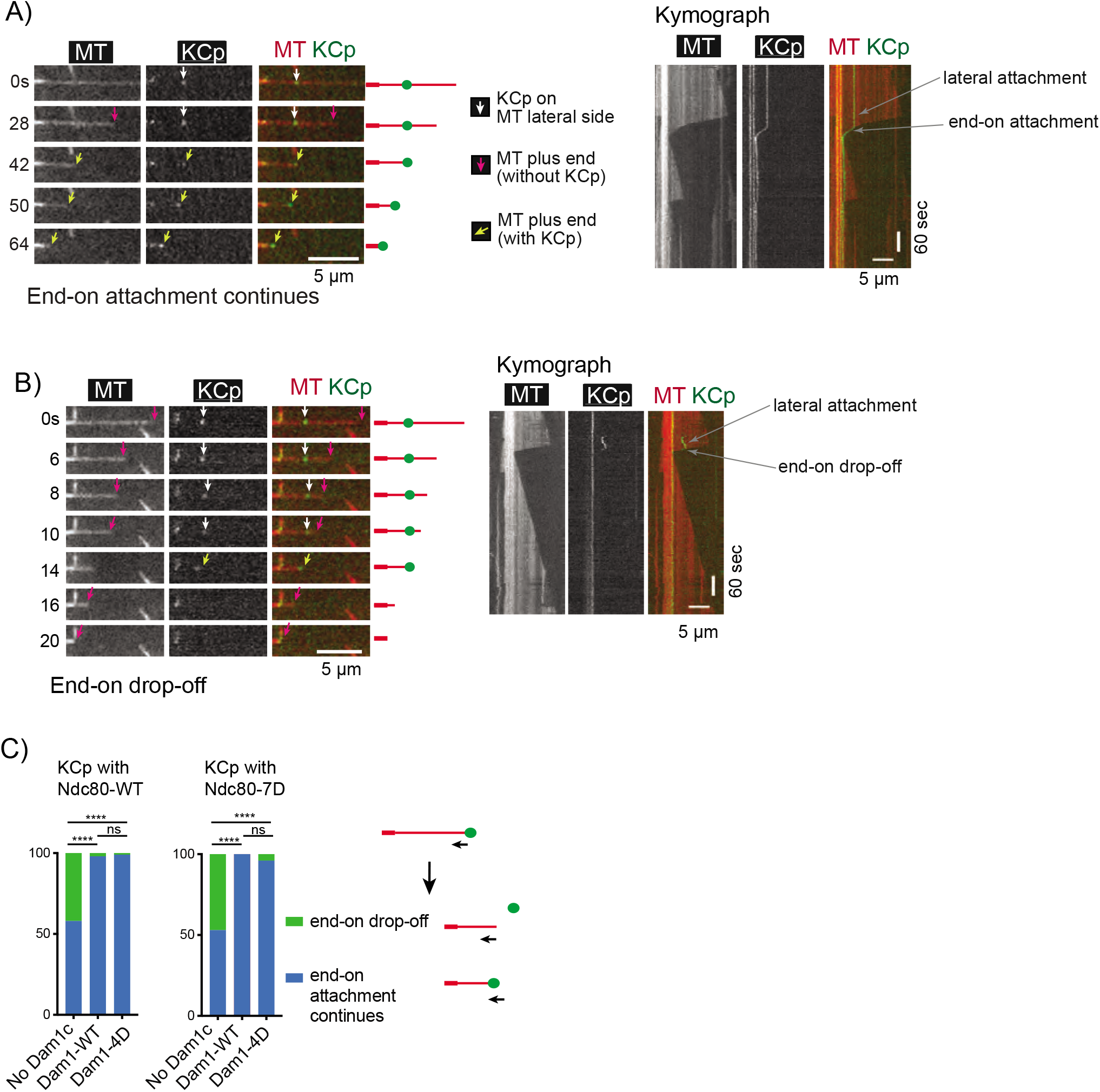
Supplemental data associated with Figure 4. A) Images in time sequence show that the KCp (purified from Dam1-depleted cells) is attached to the lateral side of a MT (white arrows, 0–28s) and formed the end-on attachment (yellow arrows, 42-64s) in the presence of recombinant Dam1C. Kymograph obtained for the same MT is shown on the right side. In this example, the KCp contained Ndc80C-WT, and recombinant Dam1C-WT was added to the system. B) Images in time sequence show that the KCp (purified from Dam1-depleted cells) is attached to the lateral side of a MT (white arrows, 0–10s), formed the end-on attachment (yellow arrows, 14s) and detached from the MT end (14–16s) in the absence of recombinant Dam1C. After the KCp had detached from the MT end, it was not visible by TIRF microscopy because it was no longer close to the coverslip. Kymograph obtained for the same MT is shown on the right side. In this example, the KCp contained Ndc80-7D. C) Percentage of various events observed for the KCp with Ndc80-WT (left) or Ndc80-7D (right) while the KCp tracked the end of a shrinking MT. The KCp was purified from Dam1-depleted cells. Experiments were conducted in the absence of recombinant Dam1C or in the presence of recombinant Dam1C-WT or Dam1C-4D (from left to right, n= 79, 63, 70 for the KCp with Ndc80-WT; n= 57, 50, 48 for the KCp with Ndc80-7D).

**Table S1:** Kinetochore proteins identified in purified kinetochore particles by mass spectrometry

| Protein name | Unique peptides | Mol. weight [kDa] | Sequence lengths | Q-value | Score | Sequence coverage | Intensity (iBAQ) |
| --- | --- | --- | --- | --- | --- | --- | --- |
| Ndc80 | 23 | 80.486 | 691 | 0 | 115.53 | 37.3 | 36449000 |
| Spc24 | 11 | 24.603 | 213 | 0 | 87.873 | 70.4 | 97405000 |
| Spc25 | 10 | 25.244 | 221 | 0 | 41.394 | 46.6 | 55587000 |
| Nuf2 | 8 | 52.973 | 451 | 0 | 27.51 | 24.4 | 18298000 |
| Spc105 | 19 | 104.82 | 917 | 0 | 110.14 | 25.8 | 23655000 |
| Kre28 | 8 | 44.674 | 385 | 0 | 56.648 | 26.5 | 40184000 |
| Nnf1 | 11 | 23.639 | 201 | 0 | 107.89 | 50.7 | 61039000<br>0 |
| Nsl1 | 17 | 25.416 | 216 | 0 | 207.09 | 77.8 | 47226000<br>0 |
| Mtw1 | 26 | 33.243 | 289 | 0 | 187.52 | 58.8 | 41842000<br>0 |
| Dsn1 | 9 | 65.691 | 576 | 0 | 37.745 | 22.9 | 10104000 |
| Cnn1 (CENPT) | 1 | 41.312 | 361 | 0 | 8.3498 | 5.5 | 679690 |
| Wip1 (CENPW) | 2 | 10.244 | 89 | 0 | 8.5453 | 39.3 | 1049100 |
| Okp1 (CENPQ) | 14 | 47.349 | 406 | 0 | 89.043 | 38.9 | 29199000 |
| Ame1 (CENPU) | 10 | 37.461 | 324 | 0 | 29.306 | 35.2 | 27016000 |
| Mcm21 (CENPO) | 3 | 42.97 | 368 | 0 | 2.56 | 10.6 | 15056000 |
| Ctf19 (CENPP) | 4 | 42.782 | 369 | 0 | 8.3953 | 13.6 | 7920900 |
| Mif2 (CENPC) | 14 | 62.472 | 549 | 0 | 47.672 | 27 | 40790000 |
| Nkp1 | 7 | 26.976 | 238 | 0 | 30.398 | 34.5 | 39782000 |
| Iml3 (CENPL) | 5 | 28.066 | 245 | 0 | 17.633 | 20.4 | 7922700 |
| Nkp2 | 1 | 17.862 | 153 | 0 | 6.1052 | 7.8 | 4711800 |
| Chl4 (CENPN) | 2 | 52.671 | 458 | 0 | 6.2515 | 7 | 1312700 |
| Mcm16 (CENPH) | 1 | 21.138 | 181 | 0.001674<br>1 | 1.2033 | 5 | 1627800 |
| Mcm22 (CENPK) | 2 | 27.567 | 239 | 0 | 3.0511 | 12.1 | 2709900 |
| H2B2 | 10 | 14.237 | 131;131 | 0 | 64.656 | 64.1 | 73309000<br>0 |
| H2A2 | 3 | 13.989 | 132;132 | 0 | 13.501 | 29.5 | 48168000<br>0 |
| H4 | 9 | 11.368 | 103 | 0 | 36.809 | 65 | 30438000<br>0 |
| Cse4 (CENPA) | 6 | 26.841 | 229 | 0 | 11.637 | 23.1 | 73007000 |
| Skp1 | 8 | 22.33 | 194 | 0 | 28.335 | 45.9 | 89698000 |
| Cep3 (CBF3B) | 1 | 71.357 | 608 | 0.005965<br>3 | 0.91945 | 1.6 | 396870 |
| Mad1 | 6 | 87.65 | 749 | 0 | 16.827 | 12.8 | 2212400 |
| Bub3 | 1 | 38.444 | 341 | 0 | 1.7648 | 5.3 | 745730 |
| Bub1 | 1 | 117.87 | 1021 | 0.010021 | 0.65098 | 1.1 | 0 |
| Fin1 | 3 | 33.186 | 291 | 0 | 8.4003 | 12.7 | 3210200 |
| Stu2 | 2 | 100.92 | 888 | 0 | 28.516 | 3.2 | 583010 |
| Slk19 | 1 | 95.379 | 821 | 0 | 3.3914 | 1.6 | 0 |
| Dam1 | 3 | 38.421 | 343 | 0 | 9.9438 | 13.7 | 18353000 |
| Ask1 | 4 | 32.071 | 292 | 0 | 36.05 | 29.1 | 6556500 |
| Dad3 | 2 | 10.848 | 94 | 0 | 4.6574 | 38.3 | 2579800 |
| Duo1 | 3 | 27.473 | 247 | 0 | 26.239 | 13.4 | 1215200 |
| Dad2 | 1 | 15.071 | 133 | 0 | 1.8749 | 8.3 | 0 |
| Dad4 | 1 | 8.1552 | 72 | 0 | 2.8758 | 20.8 | 0 |
| Hsk3 | 1 | 8.0881 | 69 | 0 | 7.3375 | 15.9 | 0 |
| Spc19 | 0 | 18.9007 | 165 | - | - | - | 0 |
| Spc34 | 0 | 34.0802 | 295 | - | - | - | 0 |
| Dad1 | 0 | 10.5067 | 94 | - | - | - | 0 |

Unique peptides: Number of Unique peptides identified for each protein
Molecular weight: molecular weight of the protein in kilo daltons
Sequence lengths: The amino acid length of the protein
Q-Value: a measure of false discovery rates. (Stringency of FDR (false discovery rates) cut-off)
Score: Highest Andromeda score
Sequence coverage: The % of total sequence of the protein covered by the identified peptides.
iBAQ: Intensity-based absolute quantification.
The proteins in the same sub-complex are highlighted in the same colour.

